# Multi-nucleation in two-cell human embryos stems from spindle and metaphase plate incoherence in the first mitosis

**DOI:** 10.1101/2025.08.20.671312

**Authors:** Gerard H. Pieper, Mansour Aboelenain, Lucy Munro, Bettina P. Mihalas, Cerys E. Currie, Deborah Taylor, Daniel M. Collins, Geraldine M. Hartshorne, Andrew D. McAinsh, Richard A. Anderson, Adele L. Marston

## Abstract

The first embryonic division in humans is highly error-prone and a source of aneuploidies. Multi-nucleation is prevalent in two-cell human embryos, with unknown cause. Here, we live-image human zygotes to elucidate the features of the first mitosis that predispose embryos to multinucleation. We show that failure to collect chromosomes into a single mass and establish a bipolar spindle during zygotic metaphase leads to severe multi-nucleation. We find that KIF10/CENP-E kinesin activity is essential to prevent the formation of multiple spindle poles and to congress chromosomes onto a metaphase plate. Furthermore, although the spindle assembly checkpoint mediates a delay in response to a highly disorganised spindle, KIF10/CENP-E-inhibited embryos ultimately undergo the first mitosis with a chaotic anaphase. Therefore, defective chromosome congression in the zygote combined with a failure to sense this error cause multi-nucleation. Remarkably, multi-nucleation can be corrected during the second mitotic division in a KIF10/CENP-E-dependent manner. We suggest that multi-nucleation may be a safeguarding mechanism to prevent chromosome loss during the highly error-prone first embryonic mitosis.

## Introduction

Mosaic aneuploidy is common feature of human embryos and originates from an erroneous first mitosis in the zygote^1,2^. Multi-nucleation is a common feature in human pre-implantation embryos in the IVF setting^3–8^. Using live-imaging techniques, observed multi-nucleation rates range from 23.2%^4^ - ∼43.7^6,7,9^ at the two-cell stage. Multi-nucleation peaks at the two-cell stage and decreases with subsequent divisions^5–7,9^, undergoing correction^9,10^. The relationship between multi-nucleation at this stage and embryo aneuploidy is unclear^9,11^. However, multi-nucleation at various stages of pre-implantation development correlates with lower rates of live-births after embryo transfer^4,6–9^. Nevertheless, multi-nucleated embryos can give rise to healthy live-births^4,6–9^, suggesting that multi-nucleation per se is not incompatible with normal subsequent development.

Several causes of multi-nucleation in zygotes have been suggested. In human zygotes, unfocussed spindle poles and a low spindle aspect ratio are correlated to multi-nucleation, though the mechanism is unclear^10^. The parallel positioning of the cleavage plane with respect to the orientation of the pronuclei and mitotic spindle is also important for preventing multi-nucleation^12^. Mouse zygotes with distant pronuclei form multiple unfused spindles, leading to multi-nucleated blastomeres^13^. Similarly, in bovine zygotes, defects in centrosome-mediated clustering of the pronuclei can contribute to both mitotic errors and micronuclei^14^. Lagging chromosomes can also contribute to the formation of micronuclei in human zygotes^1^. However, it is unclear if micronuclei are formed through different mechanisms than those generating a multi-nucleated state. Thus, a clear molecular mechanism that explains the high incidence of multi-nucleation in human zygotes is currently lacking.

Here, we have employed live-cell time-lapse imaging of human embryos to investigate the origin of multi-nucleation during the first mitotic division, and the impact of this on the second mitotic division. We find that multi-nucleation is caused by failed chromosome congression and multi-polar spindles in the error-prone first mitosis. We propose that multi-nucleation may serve as a safeguarding mechanism to retain genetic information for subsequent, higher fidelity, divisions.

## Results

### A system to study the causes of multinucleation in human embryos

Multi-nucleation is frequently observed in normally fertilized two-pronuclear (2PN) human zygotes in the clinical setting ^3–8^. However, studying this phenomenon is challenging since 2PN zygotes are the goal of treatment cycles and are largely unavailable for research. To overcome this, we reasoned that deselected one-(1PN) and three-pronuclear (3PN) zygotes, which are unsuitable for treatment, could provide an ideal model to study the origins of multi-nucleation in the first and second mitotic divisions. Importantly, the zygotic genome is largely transcriptionally silent at this stage^15^ and therefore cytoplasmic changes are expected to be minimal, despite the differences in chromosome number. Moreover, we anticipated that the greater chromosome number in 3PN zygotes could increase multi-nucleation frequency, providing greater power to study this phenomenon where material is limited.

We first quantified multi-nucleation in untreated deselected zygotes after completion of the first mitotic division. We found that ∼85% (12/14) of first mitoses resulted in some form of multi-nucleation in the daughter cells, around half of which we classified as minor multi-nucleation (2-3 nuclei/cell), with the remaining half classified as massive multi-nucleation (3+ nuclei/cell) (Fig. 1A, B). Previous work detected a higher rate of multi-nucleation in women ≥ 40 years as compared to ≤ 35 years old^9^. Our data show a similar trend, though the p-value was only 0.22, potentially due to insufficient numbers of embryos (Supplemental Fig. 1A). Therefore, deselected zygotes provide a powerful system to study the causes of multi-nucleation.

**Figure 1.**
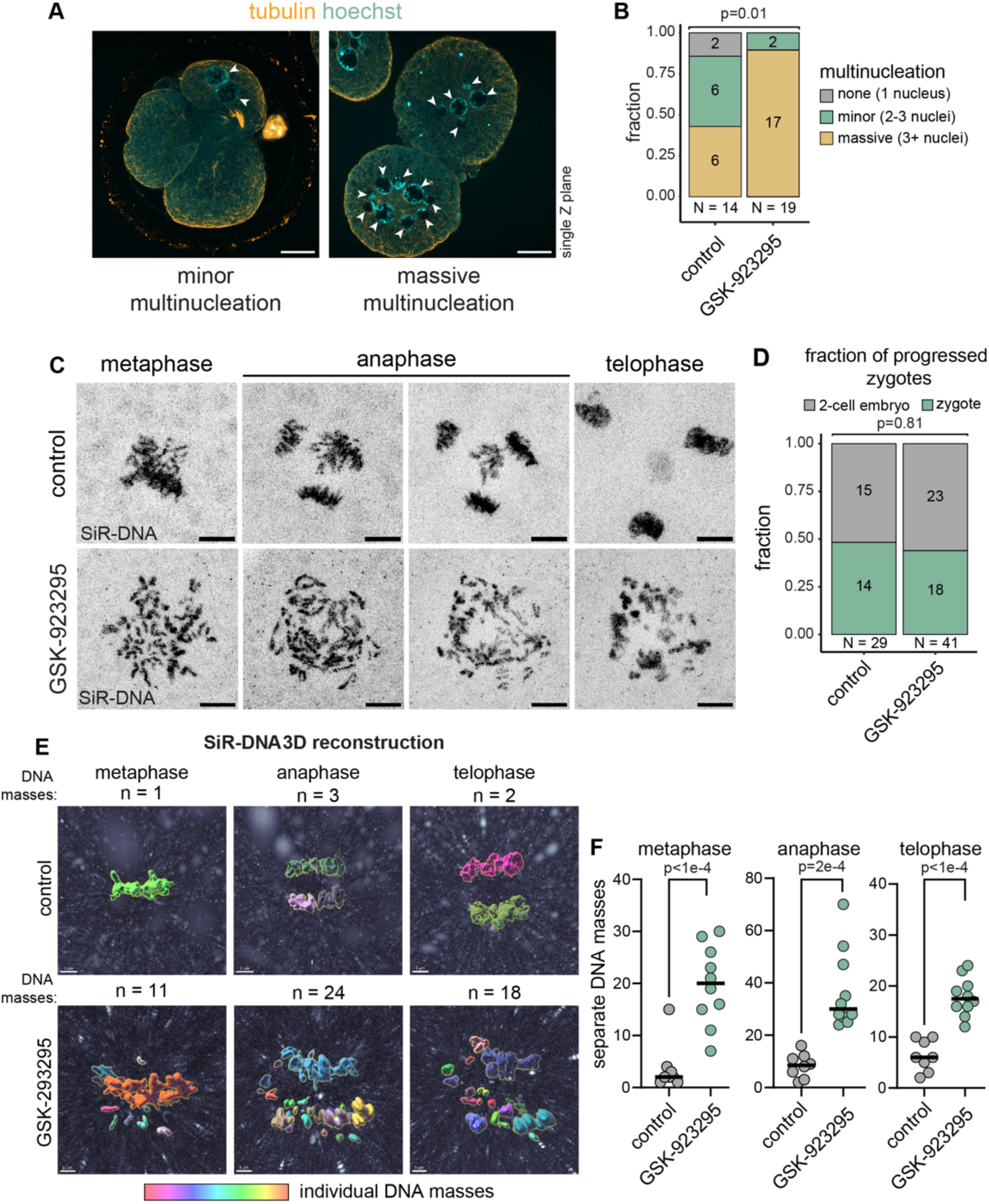
CENP-E kinesin activity prevents multi-nucleation during the first embryonic mitosis. A) Representative images from time-lapse experiments microscopy experiments of zygotes incubated with SiR-DNA and indicated drug treatments. Scale bar represents 5μm. B) Quantification of the number zygotes that progressed to embryo stage within 15-20hrs of incubation. P-value calculated with Fisher’s exact test. C) Representative 3D Imaris reconstructions of chromosomes of a control and GSK-923295-treated zygote. Individual chromosome masses are individually coloured. Scale bar represents 5μm. D) Quantification of separately identifiable DNA-masses based on 3D chromosome reconstructions in Imaris. p-value calculated by Welch’s t-test. N=8 for control, N=10 for GSK-923295 treated group. E) Representative images of minor and massively multi-nucleated human embryos. Scale bar represents 20μm. F) Quantification of multi-nucleation in human embryos after 15-20 hrs of incubation with indicated treatments. We included in this group untreated or DMSO treated zygotes which were incubated overnight in an incubator (for 15-20 hrs) to allow progression through mitosis 1 and also those that were used for live-imaging of mitosis 1 (Fig. 1-3). p-value calculated with Fisher’s exact test.

### Dispersed chromosomes cause multi-nucleation during the first embryonic mitosis

We hypothesised that multi-nucleation could be a result of a failure to segregate chromosomes into just two masses after the first mitosis, leading to each DNA mass forming a separate nucleus. We reasoned that inducing chromosome dispersal in zygotes might increase the number of chromosome masses produced during anaphase of the first mitosis. In cultured somatic cells, inhibition of the KIF10/CENP-E kinesin motor leads to polar and misaligned chromosomes^16–19^. We therefore treated zygotes with the KIF10/CENP-E inhibitor GSK-923295^20^ and followed them by live imaging as they progressed into the first mitosis (Fig. 1C). A comparable proportion of control and GSK-923295-treated zygotes, roughly half, progressed through the first mitosis (Fig. 1D). Following overnight incubation, these zygotes produced a variable number of cells, which was not different between control and GSK-923295 treatment (Supplemental Fig. 1B, C). Notably, embryos with 3 cells potentially reflect tri-polar divisions^21^.

To ask whether GSK-923295 treatment perturbs chromosome congression in metaphase human zygotes, as observed in cultured somatic cells^22,23^, we quantified the number of separately identifiable chromosome masses, based on the SiR-DNA signal. This revealed that compared to a median of 2 in control cells, GSK-923295-treated zygotes exhibited around 20 individual DNA masses in metaphase of mitosis 1 and individual chromosomes were readily observable in GSK-923295 treated embryos (Fig. 1C, E, F). This confirms that KIF10/CENP-E promotes chromosome congression in human zygotes and demonstrates that KIF10/CENP-E inhibition can induce chromosomal dispersion in human embryos. We note, however, that the observed phenotype upon KIF10/CENP-E inhibition differs substantially between human zygotes and cultured somatic cells^22,23^. In both cases, KIF10/CENP-E promotes chromosome congression into the metaphase plate. Conversely, KIF10/CENP-E inhibition in zygotes, has a more extreme effect on chromosome alignment. Importantly, the number of DNA masses increased significantly in anaphase, in both control and GSK-923295-treated groups, indicating loss of sister-chromatid cohesion and the separation of chromosome masses along the spindle (Fig. 1C, E, F). GSK-923295-induced chromosome dispersion persisted through anaphase and telophase, since the number of DNA masses was significantly higher than in control zygotes (Fig. 1F). Moreover, these DNA masses were frequently packaged into separate nuclei at telophase (Fig. 1F), which led to an increased rate of multinucleation in the GSK-923295-treated zygotes (Fig. 1A, B). We conclude that a failure to cluster chromosomes prior or following their segregation results in multi-nucleation and that the KIF10/CENP-E kinesin motor plays a major role in preventing multi-nucleation by clustering chromosomes in human zygotes.

Age of the oocyte donor did not play a role in the multi-nucleation phenotype as maternal age was similar in all treatment groups (Supplemental Fig. 1D). Our data set contains a small number of 1PN zygotes. The fraction of 1PNs and 3PNs in each treatment group was not significantly different (Supplemental Fig. 1E). Additionally, amongst control zygotes, a difference in multi-nucleation state between 1PNs and 3PNs was not detected (Supplemental Fig. 1F). We also observed frequent embryo fragmentation, which was comparable in all conditions (Supplemental Fig. 1G).

### Failed chromosome congression and a lack of spindle coherence lead to multi-nucleation during the first embryonic mitosis

Our finding that dispersed chromosomes in zygotes result in multi-nucleation after the first mitosis prompted us to investigate why chromosomes might fail to cluster in embryos. A failure of chromosomes to congress at the metaphase plate would prevent the initial clustering of chromosomes into a single mass. An alternative, not mutually exclusive possibility, is that a lack of spindle polarity pulls chromosomes in separate directions after initial clustering has been established. We predicted that, if the latter scenario is true, prolonged CENP-E inhibition would exacerbate the chromosome clustering defect.

We therefore assessed the effect of a short (30 min) CENP-E inhibition in zygotes that had undergone pro-nuclear breakdown (PNBD) or long-term GSK-923295 treatment (> 15 hrs of inhibition) of PN-stage zygotes that wer allowed to undergo mitotic entry on spindle morphology and chromosome alignment by immunofluorescence (Fig. 2A). After 30 min of inhibition, we observed that chromosomes were aligned on a small number (∼3) of spindle poles. Long term CENP-E inhibition revealed a severe loss of spindle and metaphase plate coherence (Fig. 2A). Spindles were highly multipolar with an average of 15 spindle poles and chromosomes were aligned between various pairs of spindle poles (Fig. 2A, B). This suggests that both metaphase plate and spindle coherence are lost over time upon loss of CENP-E function. We conclude that chromosome dispersal is associated both with a lack of chromosome clustering and loss of spindle pole coherence.

**Figure 2.**
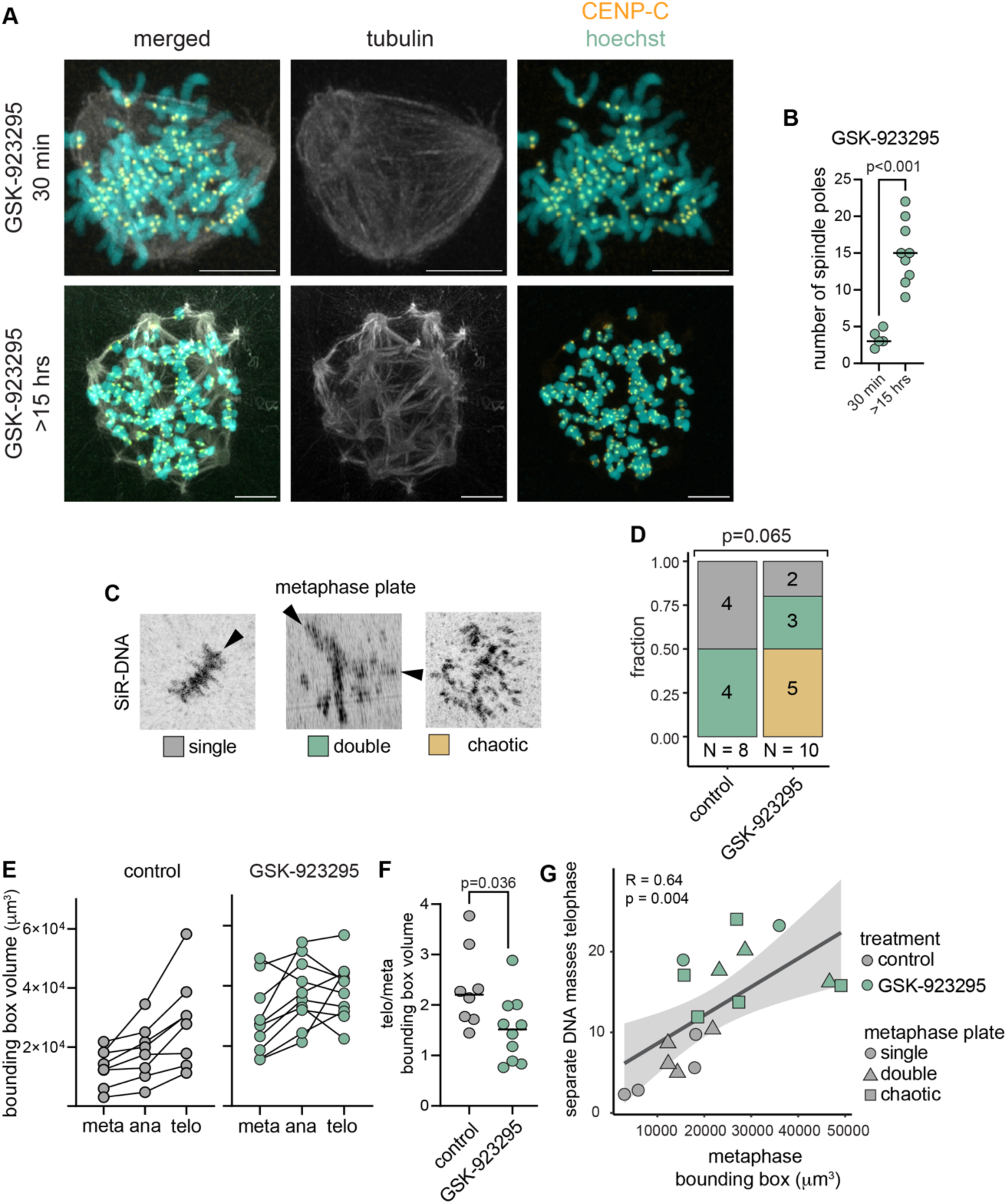
Lack of metaphase plate and spindle coherence leads to multi-nucleation during mitosis 1. A) Representative images of a metaphase zygote treated with GSK-923295 for 30 min (N=5) and of a metaphase arrested GSK-923295-treated zygote (N=9), immunostained with antibodies targeting indicated proteins and stained for DNA with Hoechst. Scale bar represents 5μm. B) Quantification of the number of spindle poles in metaphase arrested GSK-923295 zygotes. C) 3D Imaris reconstructions of metaphase “plate” appearance types. Arrowheads indicate the edges of either single or double metaphase plates. D) Quantification of metaphase plate appearance types in indicated treatment groups. P-value calculated with Fisher’s exact test. E) Quantification of bounding box volume of DNA masses at different mitotic phases and with indicated treatments. Lines represent individual cells. Bounding box volume was set as the volume of a 3D box, delimiting the outer edges of detectable SiR-DNA signal. F) Ratio of the bounding box volume of telophase over metaphase for individual zygotes for indicated treatments. Horizontal bars represent medians. P-values calculated with ANOVA and Tukey’s multiple comparisons test. G) Plot of the bounding box volume of zygotes for each treatment vs the number of nuclei being formed during telophase. Black line indicates linear regression and shaded grey area represents the confidence interval.

To determine how loss of spindle and metaphase plate coherence generates multi-nucleation we performed live-cell imaging of the first mitotic division using the live-cell DNA dye SiR-DNA (Fig. 1C, Fig. 2C). We first assessed the appearance of the metaphase plate and noted that in control zygotes, metaphase plate appearance was either “single” or “double” (Fig. 2C,D). The latter denoted a combination of two clearly definable metaphase plates, with or without the presence of additional misaligned chromosomes. The “double” plate is potentially a feature of the fact that 3PNs contain supernumerary chromosomes and centrosomes^24^. Consistently, anaphases were overwhelmingly multipolar (Supplemental Fig. 2A). In GSK-923295-treated zygotes, a new class of “chaotic” metaphase plates were observed, denoting zygotes without easily identifiable plates or with more than two plates (Fig. 2C,D), and the majority of plates were either “chaotic” or “double”. This is consistent with our observation that chromosomes are severely spread out and scattered between multi-polar spindles at metaphase (Fig. 2A).

To analyse the extent of chromosome scattering on the mitotic spindle we used bounding box analysis. The bounding box volume in this case is a measure of the volume chromosomes take up within the cell. In control zygotes, the bounding box volume increased as mitosis progressed (Fig. 2E). However, in GSK-923295 treated zygotes we observed that the bounding volume was initially higher in metaphase but did not increase to the same extent as in control zygotes. The latter was confirmed by the fact that the fold-change of the bounding box volume of telophase over metaphase in GSK-923295 treated zygotes was significantly lower compared to control cells (Fig. 2F). To assess if the metaphase plate organisation is a predictor for the number of nuclei that will be formed during telophase, we plotted the volume of the metaphase bounding box versus the number of separate DNA objects during telophase for both control and KIF10/CENP-E inhibited zygotes. We found a high correlation between metaphase bounding box size and number of nuclei formed during telophase (Fig. 2G). Control zygotes occupied the bottom left corner of graph, indicating smaller metaphase plate volumes and lower levels of multi-nucleation. All GSK-923295-treated zygotes occupied the top right corner, indicating larger metaphase bounding box volumes and higher levels of multinucleation (Fig. 2G). Therefore, both metaphase plate and spindle organisation are affected in human zygotes upon KIF10/CENP-E inhibition. During telophase, GSK-923295-treated zygotes fail to reform coherent nuclei. Instead, a large number of separate nuclei form, most likely due to the increased spread of chromosomes. Thus, the severe loss of coherence of both the metaphase plate and the mitotic spindle directly leads to multi-nucleation during telophase and subsequent cytokinesis.

### A robust spindle assembly checkpoint is active in the first embryonic mitosis

Despite chaotic metaphase plates and the presence of multiple spindle poles, surprisingly, GSK-923295-treated embryos progress into anaphase with a similar frequency to control zygotes (Fig 1D). However, KIF10/CENP-E inhibition cultured somatic cells show a spindle assembly checkpoint (SAC)-dependent arrest in metaphase^16–18^. This suggests that zygotic mitotic surveillance mechanisms are either unable to detect these errors in zygotes or are insufficiently robust to prevent cell cycle progression. This prompted us to address the efficacy of the spindle checkpoint in the first mitotic division. To determine whether GSK-923295-mediated chromosome mis-alignment causes a mitotic delay, we first measured the time from PNBD to anaphase by live imaging of SirDNA-labelled zygotes. In control zygotes, PNBD to anaphase duration was around 150 minutes, consistent with previous measurements^1^. However, KIF10/CENP-E inhibition extended this timing to ∼500 minutes, indicating that KIF10/CENP-E inhibition delays progression of mitosis 1 (Fig. 3A, B). This delay could be brought back down to control levels by additionally inhibiting the SAC kinase MPS1 with reversine^25^ (Fig. 3A, B). This suggests the delay was due to SAC activity. Similarly, KIF10/CENP-E inhibition led to a SAC-dependent delay during the second mitotic division (Fig. 3C). Due to technical reasons, we could not always track the delay to completion and therefore, this delay is likely an underestimate. We observed no correlation between either maternal donor age and PNBD-anaphase duration in control cells, nor between maternal age and the length of the GSK-923295-induced delay (Supplemental Fig. 3A, B). This suggests that the strength of the SAC in zygotes is not eroded by maternal age, which is consistent with a previous report that embryonic mitotic aneuploidy is independent of maternal age^26^. We note that a delay in the first mitosis was also observed in a minor group of control, but not reversine-treated, embryos (Fig. 3B), indicating that the SAC can also be activated in unperturbed embryos.

**Figure 3.**
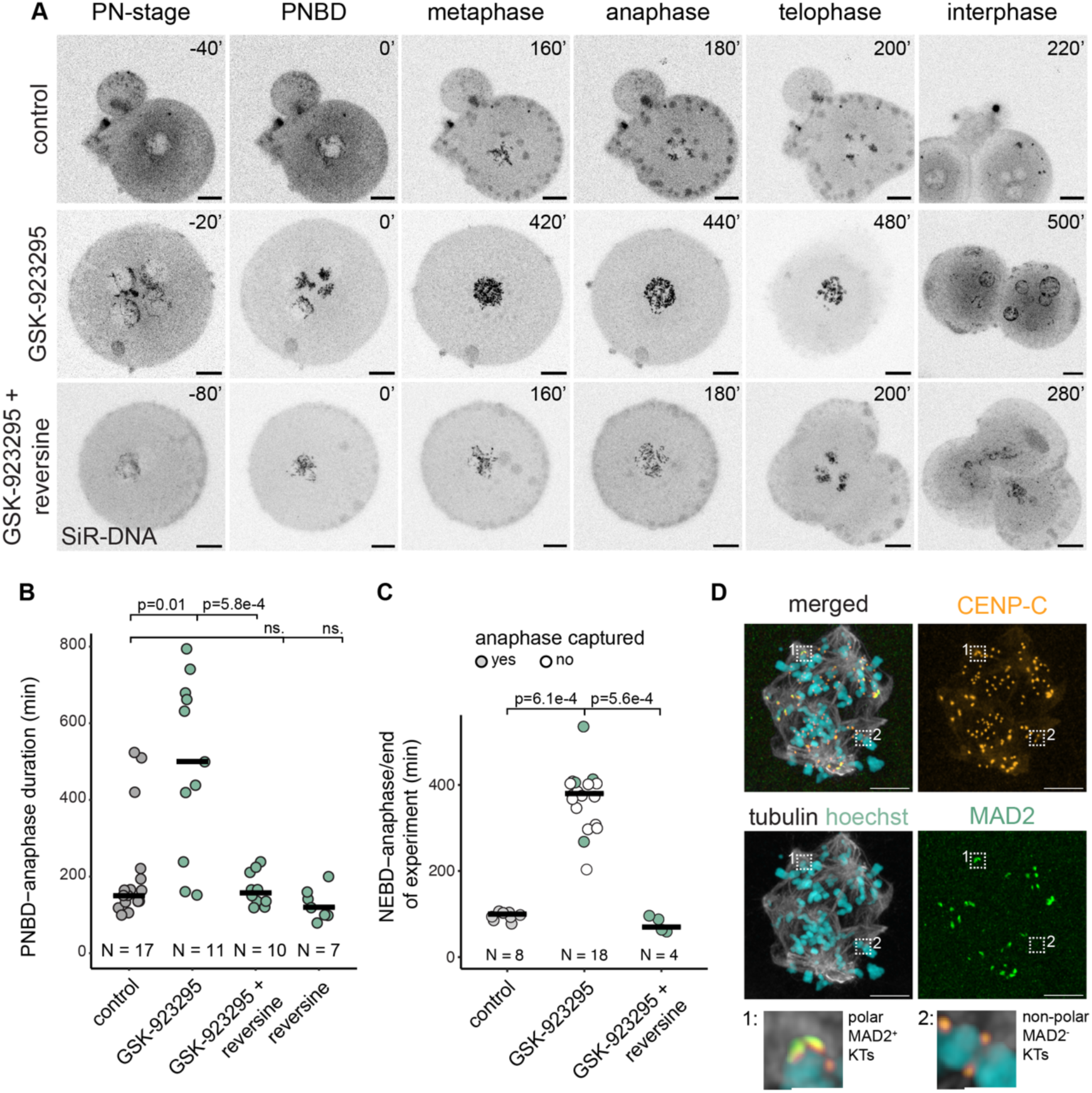
A spindle assembly checkpoint is active during embryonic mitosis 1. A) Representative time-lapse microscopy images of zygotes progressing through mitosis 1. Scale bars represent 20μm. B) Quantification of timing of the PNBD-anaphase duration of mitosis 1 for indicated treatments. P-values calculated with Kruskal-Wallis test followed by Dunn’s post-hoc test with Holm’s family-wise error correction. Only quantified zygotes that completed mitosis within the 15-20 hrs of imaging were quantified. C) Quantification of the duration of timing of NEBD-anaphase or to the end of recording for second mitosis embryos. White filled data points represent cells for which anaphase was not captured and therefore the timing refers to the duration of metaphase captured from NEBD. P-values calculated with Kruskal-Wallis test followed by Dunn’s post-hoc test with Holm’s family-wise correction. D) Microscopy image of a metaphase-arrested CENP-E inhibited (>15 hrs) stained immunostained for MAD2, CENP-C, tubulin and stained for DNA with hoechst (N=5). Scale bar represents 5μm. Inserts represent zoomed in images of MAD2 positive or negative kinetochore (KT) pairs. Scale bar represents 1μm.

Despite a clear SAC response, GSK-923395-treated zygotes eventually progressed through the first mitosis to the two-cell stage and did so with a similar frequency to control zygotes (Fig. 1D). Live imaging revealed that in both GSK-923295 and GSK-923295 + reversine treated zygotes, sister chromatid cohesion was lost abruptly, indicating a sharp transition into anaphase, despite chromosomes being in a chaotic state (Fig. 2C, Fig. 3A). As a result, GSK-923295 and GSK-923295 + reversine treated zygotes exhibited similar multinucleation (Supplemental Fig. 3C) and metaphase plate phenotypes (Supplemental Fig. 3D-F). Together, these observations suggest that zygotes can either make sufficient end-on kinetochore attachments which satisfy the checkpoint in the context of incoherent metaphase plates or that kinetochores without proper end-on microtubule attachments fail to be sensed the checkpoint.

The SAC effector, MAD2, localises to kinetochores that are not end-on attached^27^. To confirm that the SAC can sense aberrant microtubule-kinetochore attachments in KIF10/CENP-E inhibited zygotes, we incubated PN stage embryos with GSK-923295 for >15 hours prior to MAD2 immunostaining. In 5/5 embryos, MAD2 was found associated with kinetochores, though the number of MAD2-associated kinetochores varied between embryos (Fig. 3D). This indicates that unattached kinetochores retain the ability to mount a SAC signal in zygotes. These findings raise the possibility that SAC activity in zygotes, where the cytoplasm is large, requires signalling from multiple unattached kinetochores to mount a robust checkpoint arrest. We conclude that the SAC is active during the first human embryonic mitosis and can instigate a robust mitotic delay in response to a large number of erroneously attached kinetochore. However, eventually, the SAC is turned off, potentially due to insufficient signalling centres, and a coordinated, abrupt and highly multipolar anaphase ensues, which leads to multinucleation.

### Correction of multi-nucleation during the second embryonic mitosis requires KIF10/CENP-E

It has been suggested that embryos can correct or reduce the multi-nucleated state during the second embryonic division^5–7,9^. To observe this directly, we performed live-imaging of the second mitosis of deselected human embryos (Fig. 4A) and asked if KIF10/CENP-E was required for correcting multi-nucleation at this stage. Firstly, we observed that chromosomes from multinuclei always congressed into a single spindle (Fig. 4A). As reported before, we observed a reduction in multi-nucleation after the second mitosis (Fig. 4A,B). GSK-923295 and GSK-923295 + reversine treated embryos were not also able to correct the multi-nucleated state (Fig. 4A, B).

**Figure 4.**
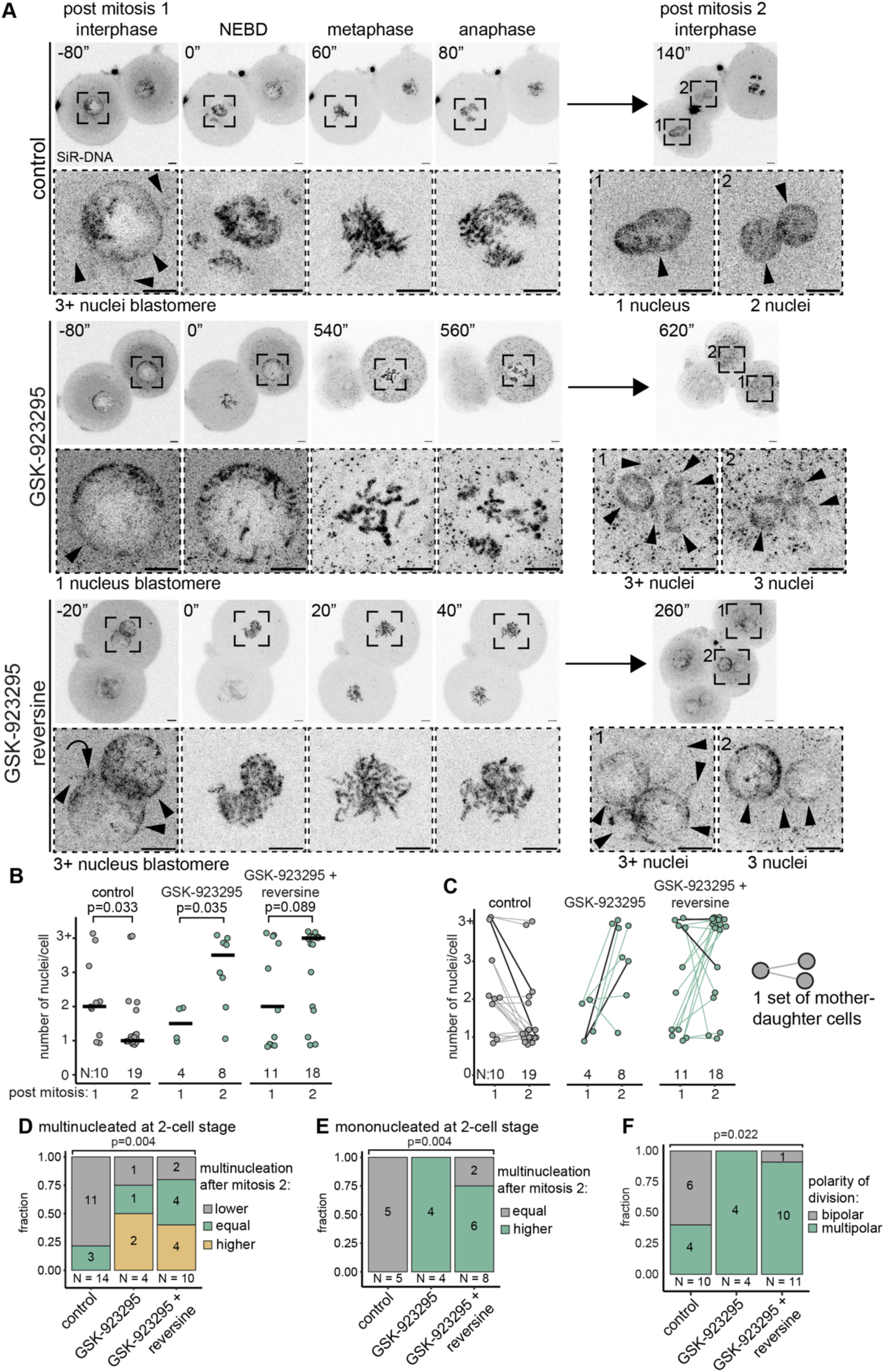
Correction of multi-nucleation during the second embryonic mitosis requires CENP-E activity. A) Representative images of time-lapse live-cell imaging of the second mitotic division of embryos incubated with SiR-DNA and indicated drugs. Scale bars represent 10μM B) Quantification of the extend of multi-nucleation in cells resulting from the second embryonic mitosis. “pre” indicates post-mitosis 1 interphase and “post” indicates post-mitosis 2 interphase. P-values calculated with Mann-Whitney U test. C) Lineage analysis of embryos during the second mitotic division. Black lines represent the embryos shown in Fig. 4A. D) Quantification of the multi-nucleation state of cells resulting from mitosis 2 which were multi-nucleated at the 2-cell stage. p-value calculated with Fisher’s exact test. E) Quantification of the multi-nucleation state of cells resulting from mitosis 2 which were mononucleated at the 2-cell stage. p-value calculated with Fisher’s exact test F) Quantification of polarity of the second mitotic division. p-value calculated with Fisher’s exact test

By lineage tracing (Fig. 4C) we found that control cells which are multi-nucleated at the two-cell stage have lower levels of multi-nucleation after the second mitosis (Fig. 4C, D). Control embryos which were mononucleated at the two-cell stage remained mononucleated after the second mitosis (Fig. 4C, E). Conversely, multi-nucleated two-cell KIF10/CENP-E inhibited and KIF10/CENP-E + SAC inhibited embryos maintained or showed high levels of multi-nucleation after the second mitosis (Fig. 4C, D). Mononucleated GSK-923295 and GSK-923295 + reversine treated embryos almost always were multi-nucleated after the second mitotic division. Unlike in mitosis 1 control zygotes, where most of the anaphases were multipolar (Supplemental Fig. 2A), during mitosis 2 a large fraction of divisions was bipolar in the control group (Fig. 4F). Interestingly, polarity of the spindle did not affect the multi-nucleation outcome in control cells (Supplemental Fig. 4B). This suggests that nucleus formation during the second mitosis might be more robust than during mitosis 1. However, KIF10/CENP-E inhibited groups were largely multipolar in both the first and second mitosis (Fig. 4F), suggesting a similar contribution of KIF10/CENP-E to metaphase plate and spindle coherence during both mitoses. In conclusion, we found that KIF10/CENP-E is required for the correction of multi-nucleation during the second mitosis. KIF10/CENP-E potentially performs this function by contributing to coherence of the metaphase plate and the spindle in the same way as during mitosis 1. In this way, KIF10/CENP-E is also required to prevent further multinucleation during the second embryonic division.

## Discussion

Here, by perturbing the organisation of chromosomes on the zygotic spindle with the KIF10/CENP-E inhibitor GSK-923295, we have shown that spindle and metaphase plate coherence prevents multi-nucleation in the first mitotic division of the human embryo. Perturbing KIF10/CENP-E activity resulted in loss of coherence of both the mitotic spindle and the metaphase plate in zygotes. This subsequently led to the formation of multiple nuclei during telophase (Fig. 5). We show that during the first embryonic mitosis a robust, but likely low sensitivity SAC is active. This eventually allows KIF10/CENP-E-inhibited zygotes to undergo anaphase with a highly disorganised spindle and chromosome organisation. During the subsequent second mitotic division, we have shown that KIF10/CENP-E is critically required for both the correction of multinucleation that arose from the first mitosis and the prevention of new multinucleation when nuclei reform at telophase of the second mitosis.

**Figure 5.**
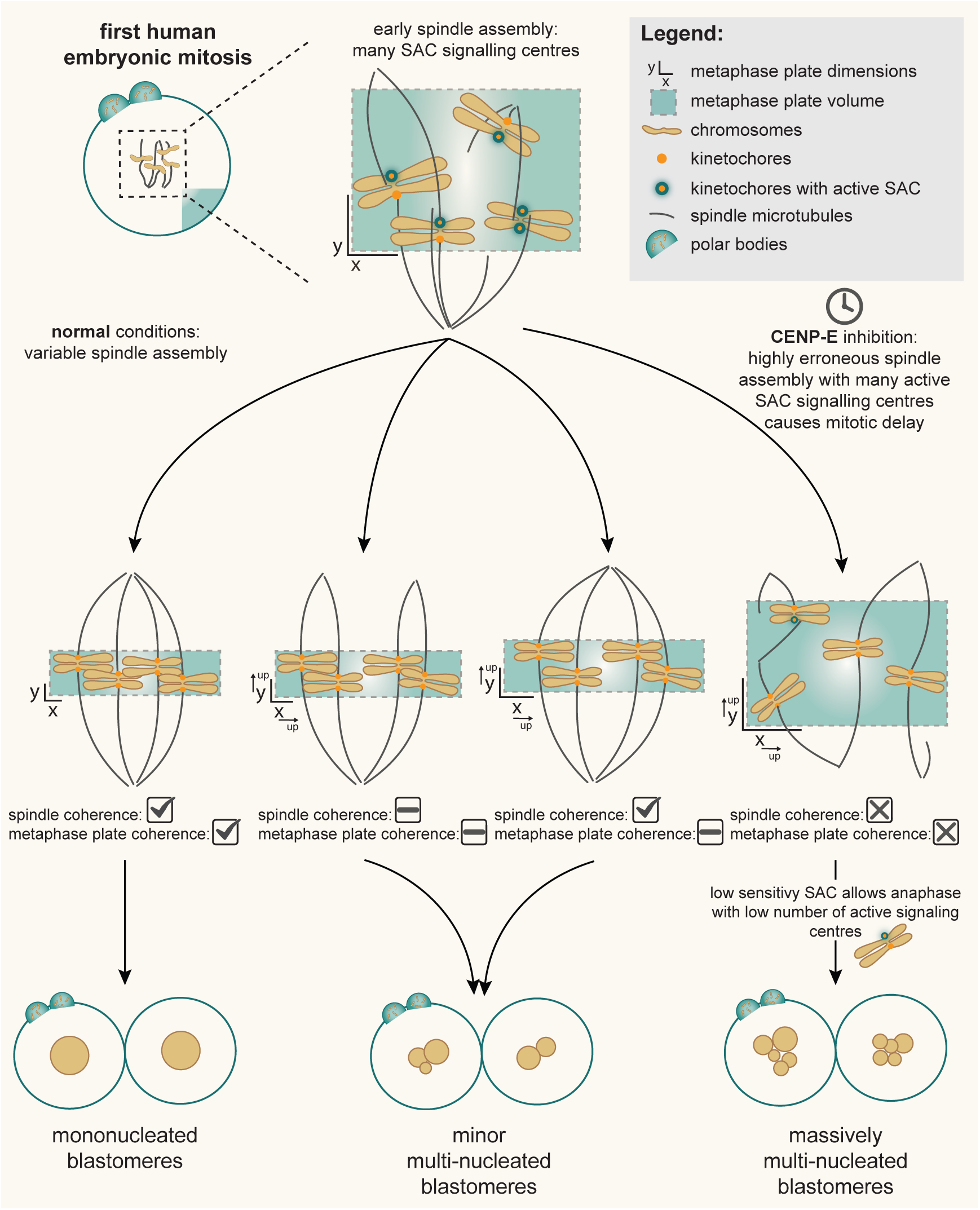
Schematic model figure.

### CENP-E is required for spindle and metaphase plate coherence in human zygotes

In mouse and human tissue culture cells and in *Xenopus* cytoplasmic egg extract, spindle organisation is not affected by KIF10/CENP-E deletion nor by antibody-mediated KIF10/CENP-E inactivation. However, several chromosomes become localised to the spindle pole, triggering a SAC arrest^16,17,19,28^. KIF10/CENP-E is critical for mouse development, as deletion of *CENP-E* resulted in an early embryonic arrest^17^. However, this is most likely due to chromosome mis-segregation throughout early development induced by polar chromosomes and weak mitotic checkpoints^17,29^. Therefore, the role of KIF10/CENP-E in spindle and metaphase plate coherence we describe here might be specific to early human development.

What could be unique about the human zygotic division that KIF10/CENP-E inhibition results in disruption of both the metaphase plate and the spindle? Though not fully investigated, there is some data to suggest that in mammalian model organisms, zygotic spindle formation does not depend on centrioles or centrosomes as in somatic cells. During mouse early development, fully developed centrosomes are only detected from the blastocyst stage^30^ and mouse zygotic spindle microtubules appear to be generated from chromosomes^13^. Furthermore, in bovine zygotes centrioles do not make a major contribution to spindle formation^31^. Human sperm-inherited centrioles can be incorporated into spindles^10,24,32^. However, whether they contribute to spindle formation, or are just loosely associated with the mitotic spindle is currently unknown. Therefore, human zygotic spindle assembly could potentially be mediated through a chromosome-based mechanism, similar to the formation of the meiotic spindle in human oocytes^33^. We speculate that in the absence of centrosome-based spindle organisation, KIF10/CENP-E might be involved in the organisation of chromosome-derived spindle microtubules into a bipolar spindle and chromosomes into a coherent metaphase plate. Alternatively, KIF10/CENP-E could be involved in establishing end-on microtubule-kinetochore attachments, which then contributes to the formation of a bipolar spindle. Similarly, in mouse oocytes, stable microtubule-kinetochore interactions are required for sorting MTOCs to the spindle poles and for spindle elongation^34^.

### Multi-nucleation is a result of loss of spindle and metaphase plate coherence

We have shown that a sharp increase in multi-nucleation is a direct result of perturbations of the coherence of the spindle and metaphase plate by interfering with KIF10/CENP-E function (Fig. 5). How can this data explain the high frequency of multi-nucleation observed in deselected and clinical-grade embryos? Firstly, for 1PN and 3PN zygotes, we show here that deselected embryos exhibit a higher rate of multi-nucleation than reported in clinical-grade embryos^3–8^. We speculate that this could be correlated to the fact that 1PNs and 3PNs have a higher incidence of multipolar spindles^1,24^ than 2PN zygotes. 1PN and 3PN zygotes can also carry an abnormal number of chromosomes, which potentially affects the organisation of the metaphase plate (Fig. 2). For clinical embryos, data suggest spindle abnormalities are a common feature in human embryos. In fact, spindle abnormalities have been detected in various stages of human pre-implantation development^35^. Furthermore, loss of spindle pole focussing is correlated to increased rates of multi-nucleation in 2PN embryos^10^. Finally, microscopy and sequencing-based methods have identified the occurrence of tripolar divisions in human embryos^1,21^. Our recent preprint suggests that zygotic metaphase plate size can be reduced by treatment with the kinesin KIF2C/MCAK agonist UMK57^2^. UMK57 treatment also led to a reduction in abnormal nuclear phenotypes such as micronuclei and a reduction in aneuploidy^2^. It will be interesting in the future to determine whether reduction of multi-nucleation is a route to improving chromosome segregation outcomes.

### A robust, but likely low sensitivity SAC is active during the first embryonic mitosis

Previous reports have speculated that the high occurrence of chromosome segregation errors in human zygotes suggests that the SAC might be dysfunctional at this stage of development^1^. Here, for the first time, we provide direct evidence that a robust SAC is active in human embryos at least during the first embryonic mitosis. Our live-imaging data shows that the zygotic SAC can delay anaphase in the face of a highly disorganised spindle in KIF10/CENP-E inhibited zygotes, which potentially exhibits many not end-on attached kinetochores. We have also shown that MAD2 is localised to metaphase kinetochores in KIF10/CENP-E inhibited zygotes. However, equal numbers of control and CENP-E inhibited zygotes do progress past mitosis 1. Indeed, our imaging data shows an instantaneous anaphase in all KIF10/CENP-E inhibited zygotes, with clear sister-chromatid separation. This precludes mitotic slippage as the underlying mechanism and instead suggests that the SAC is either silenced on all kinetochores due to the establishment of end-on microtubule-kinetochore attachments, or that the level of incorrect attachments drops below a certain detection threshold. The latter would suggest that the zygotic SAC is generally robust once a large enough number of signalling centres becomes activated. However, it might not be sensitive enough to detect small numbers of errors. Thus, low sensitivity of the checkpoint might contribute to the genesis of multinucleation in the face of a highly disorganised spindle and metaphase plate organisation. This might be a feature common to mammalian embryos as in mouse blastocysts the SAC also does not prevent anaphase in the presence of small numbers of polar chromosomes, despite MAD1 enrichment on the polar-localised kinetochore^29^. Therefore, low sensitivity of the SAC in human zygotes might contribute to the error-prone nature of human zygote mitosis and the high levels of aneuploidy observed in naturally conceived and clinical human embryos^36–38^.

### Is multi-nucleation a fail-safe mechanism in human zygotes?

We speculate that multi-nucleation might be a human zygote-specific mechanism to prevent loss of chromosomes in the face of erroneous spindle and metaphase plate formation in a large cellular volume. Multi-nucleation appears to be tolerated to some extent in early embryos. Indeed, in all multi-nucleated second divisions we observed, chromosomes from all nuclei were captured on a single metaphase plate (Fig. 4). In control zygotes, this would then result mostly in correction of the multinucleated state. A priority for future work is to determine whether multi-nucleation is correlated to levels of aneuploidy and whether reduction of multi-nucleation is sufficient to improve embryo quality.

### Limitations

Human zygotes, even deselected zygotes, are very scarce research material and depend on donations from couples undergoing IVF treatment, and therefore some experiments have a low sample size. Our findings also rely on observations with abnormally fertilised embryos (1PN and 3PN), with the assumption that 2PN embryos, which frequently lead to multi-nucleation in the clinical setting, will show similar behaviour. Nevertheless, the phenotypes reported here after KIF10/CENP-E inhibition show high penetrance and lead to robust conclusions.

## Materials and methods

### Reagents or resource tables

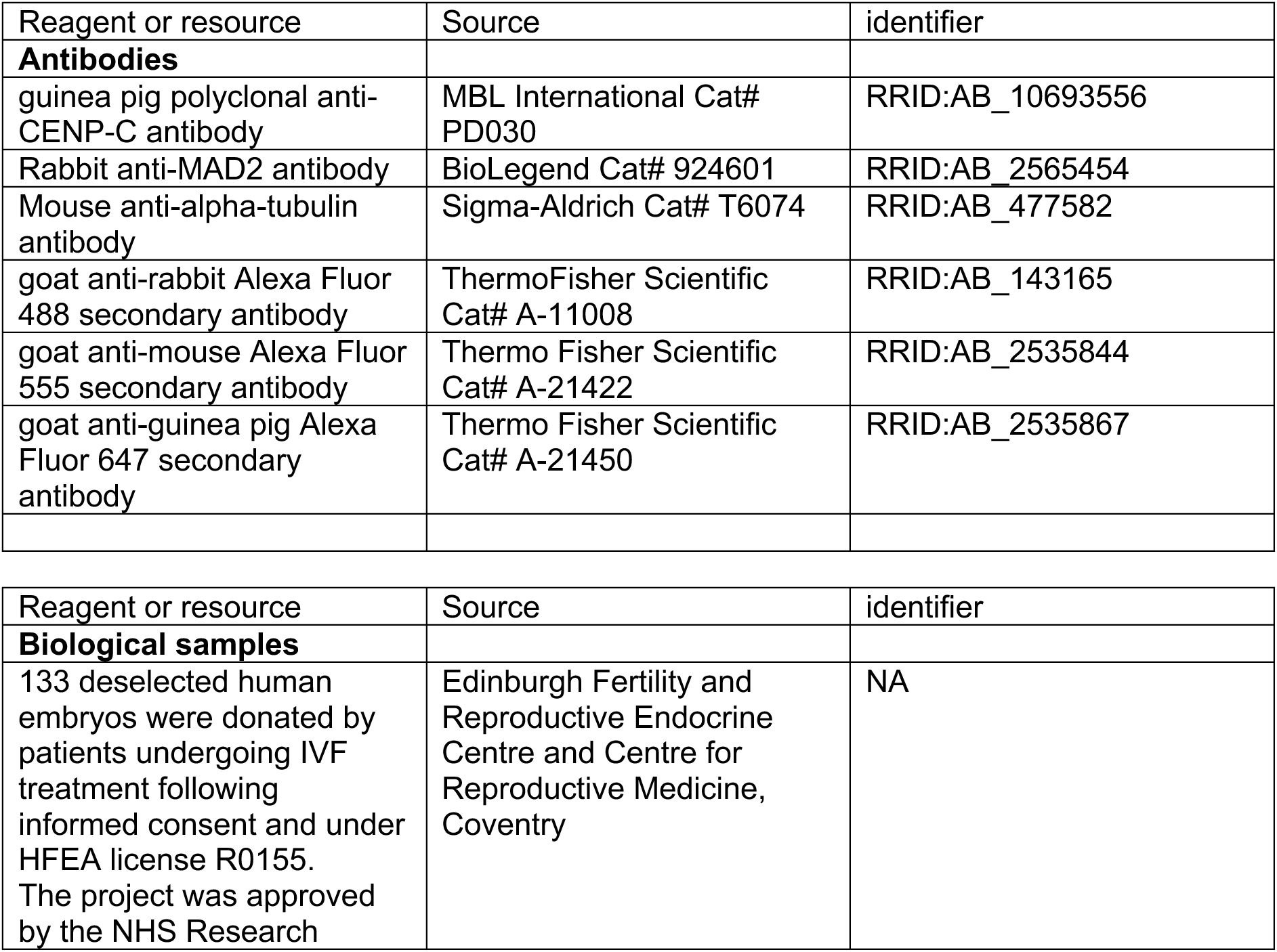
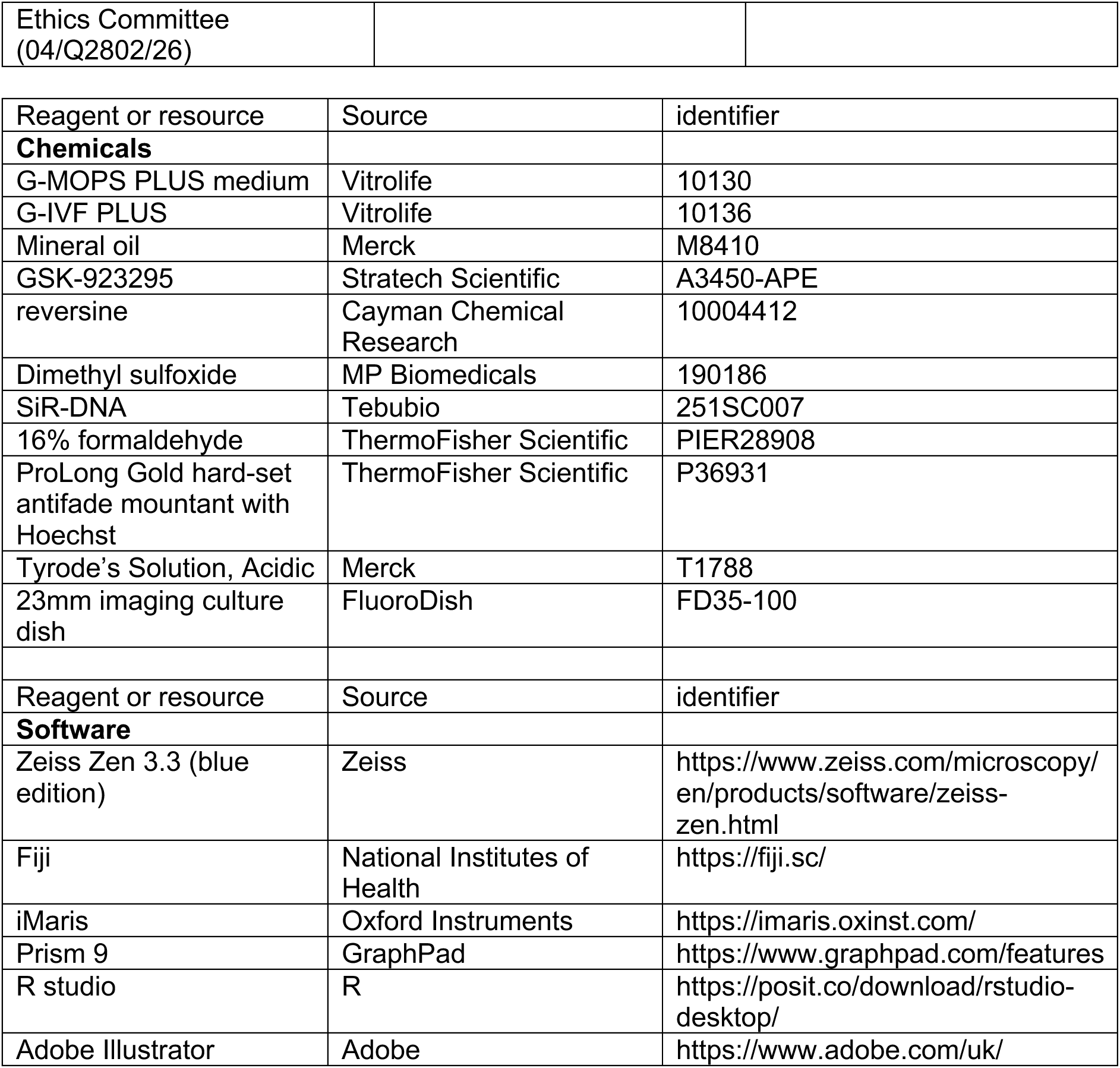

### Experimental details

#### Donation of human embryos to research

The NHS Research Ethics Committee approved this research project (Indicators of Oocyte and Embryo Development, 04/Q2802/26) and all work was conducted under a Research Licence from the Human Fertilisation and Embryology Authority (HFEA; R0155; Indicators of Oocyte and Embryo Development). Informed consent for donation of oocytes and embryos to research was provided by couples undergoing in vitro fertilisation (IVF) at the Edinburgh Fertility Centre and Department of Reproductive Endocrinology (EFC&DRE) at the Royal Infirmary of Edinburgh (NHS Lothian) or the Centre for Reproductive Medicine (CRM) at the University Hospitals Coventry and Warwickshire NHS Trust. Donations were optional and did not affect the treatment received. Couples were aware of the purpose of the research and were not provided compensation. All 1PN or 3PN zygotes were considered unsuitable for treatment and would have otherwise been disposed of.

For material collected at the EFC&DRE: Ovarian stimulation was induced using gonadotropins according to standard clinical protocols, either GnRH agonist or antagonist regimens. Oocytes were cultured and inseminated in the Vitrolife media suite at 37dC and 6% CO2. 17±1 hrs after insemination, a fertilisation check was performed by a clinical embryologist. Misfertilised (1PN or 3PN) zygotes were identified and collected from the clinic by licensed researchers 3-5 hours after the fertilisation check. Cells were transported in G-MOPS PLUS medium (Vitrolife, #10130) at 37°C in a portable incubator (K Systems) for approximately 15 min to the HFEA licensed research laboratory.

For material collected at the CRM: Ovarian stimulation was induced using FSH according to standard clinical protocols. Egg share oocytes were cultured in Origio media and inseminated by intracytoplasmic sperm injection (ICSI) using donated sperm as described previously^1^. Deselected zygotes from consenting patients undergoing routine IVF or ICSI were identified as misfertilised at the clinical fertilisation check around 17 hours after insemination, at which point they were made available to researchers. Thereafter zygotes were handled as described for material collected at the EFREC, but using Origio suite of media (Cooper Surgical).

No oocytes were vitrified or thawed in this study.

#### Embryo culture and treatments

Zygotes were cultured in G-IVF PLUS (Vitrolife, #10136) under mineral oil (Merck, # 8042-47-5) at 37°C in 5% CO_2_, 6% O_2_ and 89% N.

Upon arrival in the research lab, the stage of the embryo was confirmed by assessing the presence or absence of pronuclei (PN-stage or first mitosis respectively) or assessing the cell number.

PN-stage zygotes were incubated with either 1μM GSK-923295, 500nM reversine, a combination of the two, an appropriate DMSO control, or left untreated for 15-20 hrs. For live imaging experiments, the zona pellucida of PN-stage zygotes or two cell embryos was removed with acid Tyrode’s solution and embryos were incubated with 500nM SiR-DNA in addition to the above-mentioned treatment regimes.

#### Live-cell imaging

Cells were mounted on a FluoroDish imaging dish, covered with oil and mounted on a Zeiss LSM 980 Airyscan microscope. Cells were imaged with a 647nm laser at 1% laser power to detect SiR-DNA every 15 or 20 min with 2μm optical sections spanning the whole oocyte.

#### Immunofluorescence

Cells were first briefly washed in PHEM buffer (60 mM PIPES, 25 mM HEPES, 10 mM EGTA, 4 mM MgSO4.7H2O, pH 6.9) with 0.25% Triton X-100 at 37dC. Cells were then fixed with PHEM buffer also containing 0.25% Triton X-100 and 4% formaldehyde for 30 minutes at room temperature. Then, cells were further permeabilised for 15 min with PBS containing 0.25% Triton X-100. Cells were then washed with PBS containing 0.05% Tween-20 and stored in this buffer for up to three weeks before immunostaining.

For immunostaining, cells were first blocked with blocking solution (3% BSA and 0.05% Tween-20 in PBS) for 1hr at room temperature. Cells were then incubated with primary antibodies (anti-MAD2 1:200 from stock, anti-CENP-C 1:200 from stock, anti-tubulin 1:500 from stock) in blocking solution overnight at 4dC. Primary antibodies were washed out at room temperature in three washing steps with the final step lasting 1hr in 1% BSA and 0.05% Tween-20 in PBS. Cells were transferred to secondary antibodies at room temperature (all 1:500 from stock), which were then washed out as above. Cells were immobilised in a resin drop of ProLong Gold hard-set antifade mountant with NucBlue (Hoechst) mixed 1:1 with PBS on a Fluorodish, which was allowed to harden overnight at room temperature in the dark. Cells were then stored at 4dC and imaged within two weeks.

#### Super resolution imaging

Fixed samples were imaged using a Zeiss LSM 980 Airyscan microscope equipped with an Airyscan 2 detector and a Plan-APO (63x/1.4 NA) oil objective (Zeiss UK, Cambridge). 0.15uM optical sections were used to image entire spindles. Samples were imaged with 405nm, 488nm, 561nm and 639nm lasers to detect Hoechst, alexa-488, alexa-555 and alexa-647 respectively.

#### Image processing and data analysis

To perform deconvolution and pixel reassignment of Airyscan images, files were processed with 3D Airyscan Processing using the Auto Filter and Medium Strength functions in the Zeiss Zen 3.3 (blue edition) to generate 120 nm resolution images. For representative images of live-cell imaging experiments, Z-sections encompassing chromosomes or nuclei only were prepared extracted using Zen software for better signal/noise ratio of the final images. Representative images were prepared by making MAX Z-projections using Fiji software (National Institutes of Health).

#### Quantification and statistical analysis

For chromosome tracking and bounding box analysis of Figs 1 and 2 iMaris software (Oxford instruments) was used. The surface function was used to create 3D representation of DNA. Chromosomes were identified based on the SiR-DNA signal. Because SiR-DNA signal was highly variable between experiments, signal thresholds were manually adjusted between samples. Separate DNA objects were counted this way for each phase of mitosis. Then, all DNA objects were unified and bounding box parameters were extracted. We defined the 3D bounding box as the volume taken up by a box if it were drawn around the outer perimeters of all the DNA on the spindle.

Blebbing status was defined as follows: Minor) <5 small blebs or 1 larger bleb. Major) more than minor, mostly included large numbers of small blebs and/or very large anucleate blebs.

Spindle polarity during live-imaging was classified as either “bipolar” (segregation of chromosomes along 1 axis) or “multipolar” (segregation of chromosomes along more than 1 axis).

Statistical analysis for comparisons between conditions or mitotic stages was performed in either Graphpad Prism 9 software or R software. For each experiment, data was assessed for normality, sample size and variance between treatments. Then, appropriate statistical tests were performed as indicated in figure legends. Non-significant values were denoted as “ns.”. Legends and graphs were generated using either Graphpad Prism 9 software or with R software. Linear correlations and corresponding graphs were performed in and generated with R. Lineage tracing graph of Fig. 4C was generated in R.

## Acknowledgements

The authors wish to thank all patients who donated material allowing this research to take place as well as the clinical teams whose work supports both the patients and the research programmes. We thank Aleksandra Byrska and all Edinburgh and Warwick colleagues in the Eggs & Embryos research group for helpful discussions. We acknowledge Norma Forson for consenting all Edinburgh couples. We gratefully acknowledge David Kelly and Toni McHugh, the former Wellcome Centre Optical Imaging Laboratory (COIL) and the Discovery Research Platform for Hidden Cell Biology Light Microscopy Core for microscopy support. This work was funded by Wellcome through a Sir Henry Wellcome Fellowship to G.H.P. (222810), a Wellcome Collaborator Award (215625) (G.H.P., M.A., L.M., B.P.M., C.E.C., G.M.H., R.A.A., A.D.M., and A.L.M.), a Wellcome Investigator award to A.L.M (220780) (A.L.M., B.P.M., G.H.P., M.A., and L.M.), core funding for the Wellcome Centre for Cell Biology (203149) (G.H.P., M.A., L.M., B.P.M., and A.L.M.) and the Discovery Research Platform for Hidden Cell Biology (226791).

## Author contributions

Conceptualisation: G.H.P., A.L.M., A.D.M.; methodology: G.H.P., M.A., L.M., B.P.M., C.E.C.; formal analysis: G.H.P.; investigation: G.H.P., M.A., L.M., B.P.M., C.E.C.; resources: D.T., D.M.C., G.M.H., R.A.A.: writing-original draft: G.H.P., A.L.M.; writing-review and editing: G.H.P., A.L.M., A.D.M., M.A., G.M.H., D.M.C., R.A.A., , visualisation: G.H.P.; supervision: A.D.M., A.L.M.; project administration: G.H.P., R.A.A., A.L.M., A.D.M.; funding acquisition: A.L.M., A.D.M., G.H.P., R.A.A., G.M.H.

