## Supplemental Data for "Multi-nucleation in two-cell human embryos stems from spindle and metaphase plate incoherence in the first mitosis"

### Supplemental Figure 1

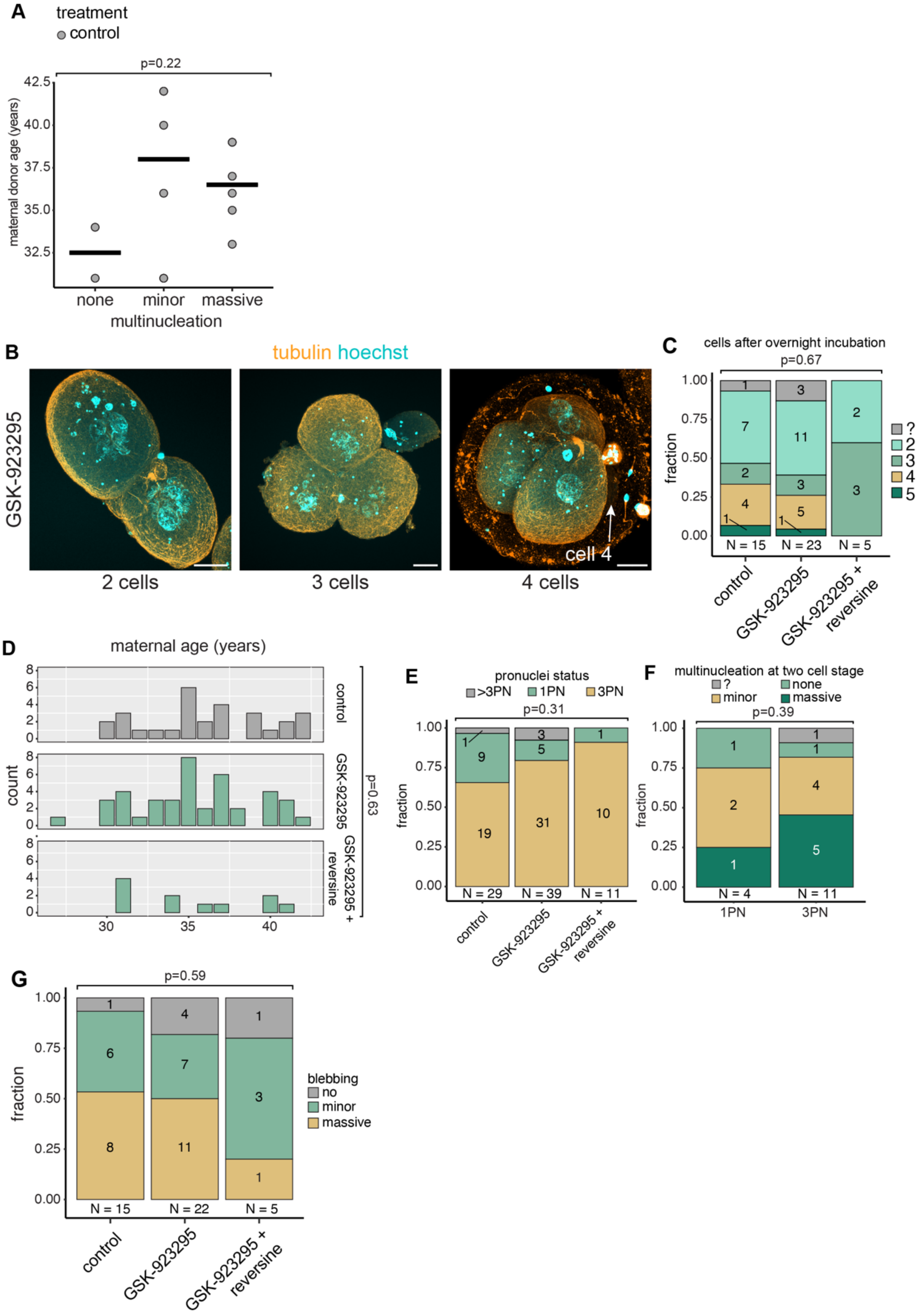


**Supplemental Figure 1. GSK-923295 treatment does not affect embryo cell number.**

A) Quantification of the maternal donor age of control zygotes with indicated phenotypes. p-value calculated with ANOVA.

B) Representative images of multicellular embryos obtained after incubating zygotes for 15-20 hrs. Scale bar represents 20µm.

C) Quantification of the number of embryo cells after 15-20 hrs of incubation. p-value calculated with Fisher's exact test.

D) Quantification of the maternal donor age of donated zygotes in each of the indicated treatment groups. P-value calculated by ANOVA.

E) Quantification of the PN-status of zygotes in the different treatment groups. p-value calculated with Fisher's exact test.

F) Quantification of the level of multi-nucleation in 1PN and 3PN zygotes. p-value calculated with Fisher's exact test.

G) Quantification of the extend of blebbing in embryos obtained after 15-20hrs of incubation. p-value calculated with Fisher's exact test.

**Supplemental Figure 2**

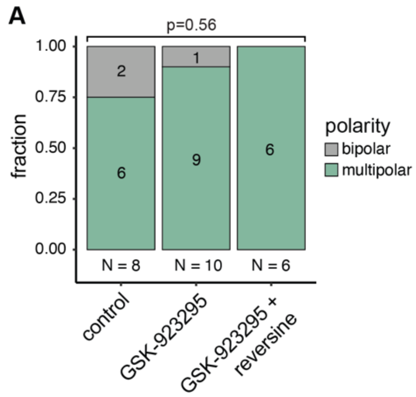


**Supplemental Figure 2. Multipolar divisions are abundant in deselected zygotes.**
A) Quantification of bipolar and multipolar divisions during mitosis 1 for the indicated treatments. p-value calculated with Fisher’s exact test.

**Supplemental Figure 3**

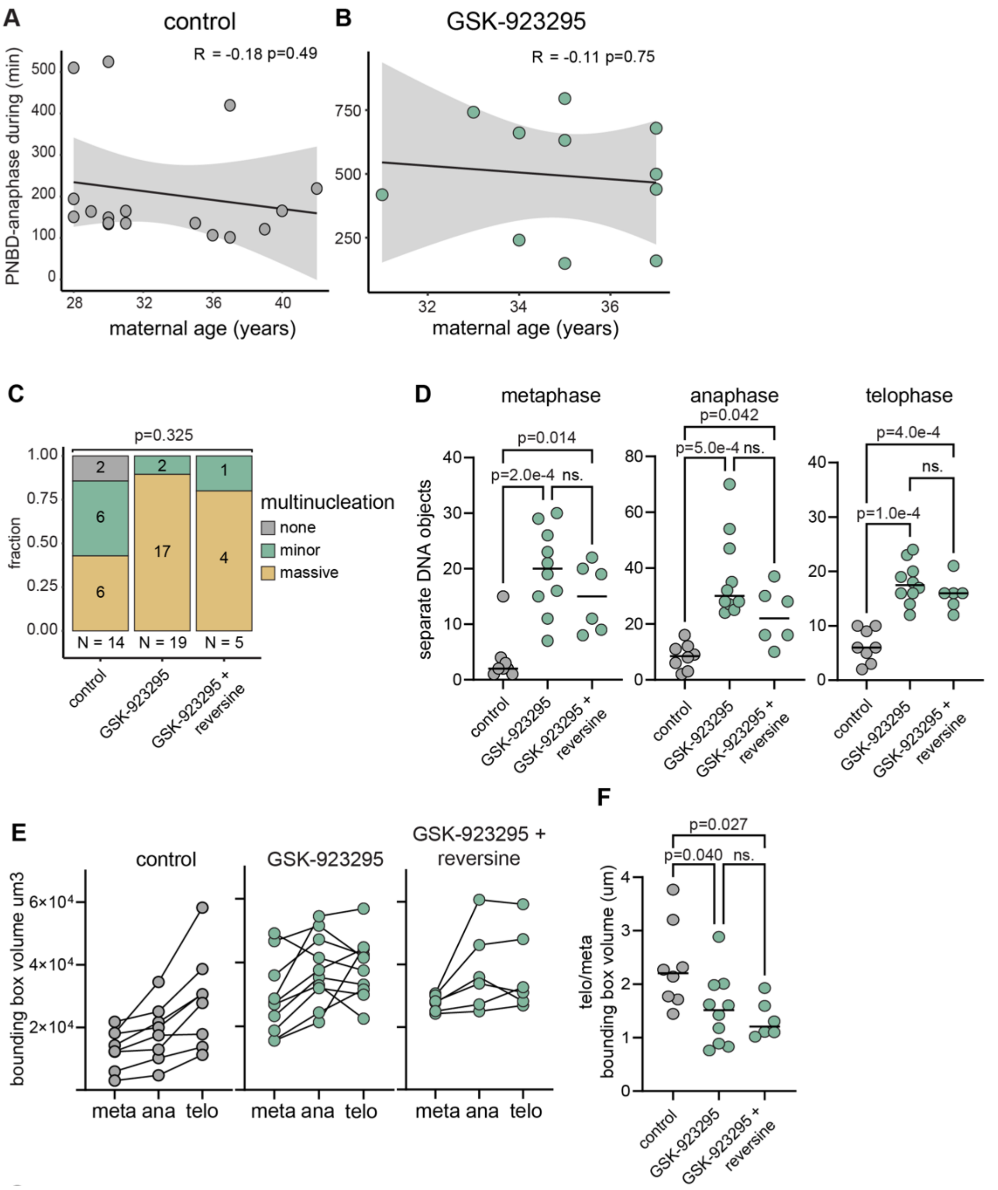


**Supplemental Figure 3. GSK-923295 + reversine treated zygotes behave similarly to GSK-923295 treated zygotes.**

A) Correlation between maternal donor age and PNBD-anaphase duration in control zygotes.

B) Correlation between maternal donor and PNBD-anaphase duration in GSK-923295 treated zygotes

A,B) Grey line represents linear data fit, shaded region represents confidence interval. Correlation coefficient (R) and p-value calculated with Pearson's method.

C) Quantification of multi-nucleation in human embryos after 15-20 hrs of incubation with indicated treatments. We included in this group untreated or DMSO treated zygotes which were incubated overnight in an incubator (for 15-20 hrs) to allow progression through mitosis 1 and also those that were used for live-imaging of mitosis 1 (Fig. 1-3). Note that control and GSK-923295 data are replicated from Fig. 1F. p-value calculated with Fisher's exact test.

D) Quantification of separately identifiable DNA-masses based on 3D chromosome reconstructions in Imaris. Note that control and GSK-923295 data are replicated from Fig. 1D. p-value calculated with Kruskal-Wallis test.

E) Quantification of bounding box volume of DNA masses at different mitotic phases and with indicated treatments. Lines represent individual cells. Bounding box volume was set as the volume of a 3D box, delimiting the outer edges of detectable SiR-DNA signal. Note that control and GSK-923295 data are replicated from Fig. 2D.

F) Ratio of the bounding box volume of telophase over metaphase for individual zygotes for indicated treatments. Horizontal bars represent medians. P-values calculated with ANOVA and Tukey's multiple comparisons test. Note that control and GSK-923295 data are replicated from Fig. 2E.

**Supplemental Figure 4**

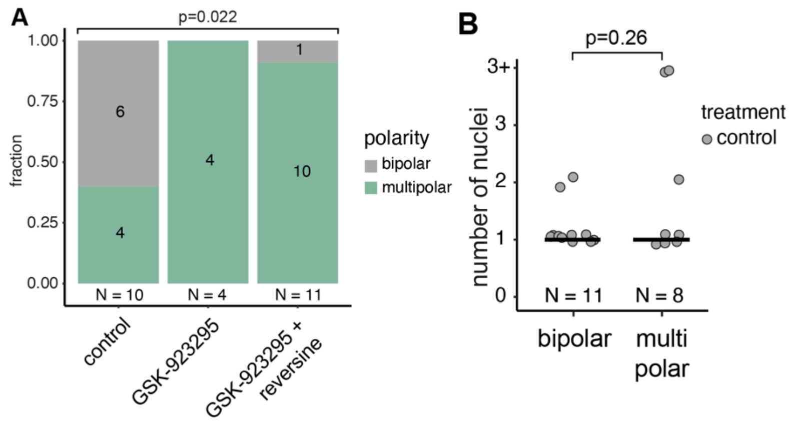


**Supplemental Figure 4. GSK-923295 treatment induces multipolar divisions during the second embryonic mitosis.**

A) Quantification of spindle polarity during the second mitotic division. p-value calculated with Fisher's exact test.

B) Quantification of the multi-nucleation state of cells resulting from bipolar or multipolar second mitotic divisions. P-values calculated with Mann-Whitney U test.
